# AFM-based nanoscale characterization of physical interaction within hematopoietic stem cells niche at single-cell level

**DOI:** 10.1101/2025.01.03.627171

**Authors:** Andrzej Kubiak, Natalia Bryniarska-Kubiak, Mehmet Eren, Kacper Kowalski, Kinga Gawlińska, Patrycja Kwiecińska, Martine Biarnes-Pelicot, Marie Dessard, Jana El Husseiny, Ti-Thien N-Guyen, Pawel Kożuch, Olga Lis, Marta Targosz-Korecka, Pierre-Henri Puech, Krzysztof Szade

## Abstract

Hematopoietic stem cells (HSCs) produce all blood cells throughout the lifespan of the organism. HSC requires a specialized bone marrow microenvironment, known as the niche, for proper differentiation and self-renewal. While several molecular and cellular elements of the niche are described, the precise understanding of the mechanobiology underlying HSC adhesion to the niche remains poorly understood. Here, we aim to characterize physical interactions and adhesion within the hematopoietic niche by combining cell sorting, surface functionalization, and atomic force microscopy.

Using this approach, we quantified and compared the adhesion of bone marrow (BM) mesenchymal stromal cells (MSCs) to different extracellular proteins, as well as the adhesion of HSCs to MSCs. We observed that MSCs adhere with the greatest force to fibronectin in an Arg-Gly-As (RGD) motif mediated and actin cytoskeleton-dependent manner. Additionally, we showed that HSCs strongly adhere to MSC within 30 seconds and that the binding is RGD-independent.

In conclusion, we demonstrated how to implement atomic force microscopy to measure the physiological interactions within the HSCs niche in a direct, specific, and quantitative way. This approach provides a more comprehensive and precise characterization of the biology of the HSC niche compared to previously used methods. Furthermore, it highlights the complexity of studying extremely small structures and rare cell populations in biophysical experiments.

## Introduction

Long-term hematopoietic stem cells (LT-HSCs) play a pivotal role in the hematopoietic system. LT-HSCs are the exclusive population capable of producing all blood cells types and self-renewing their own population. LT-HSCs reside within a specific bone marrow (BM) micro-environment known as the niche. The niche is essential for regulating LT-HSC functions, including proliferation, migration, and differentiation^1,2^. The regenerative potential of LT-HSCs warrants the therapeutic potential of BM transplantation. After transplantation, LT-HSCs home to specific BM niches and drive hematopoiesis of the transplant recipient^3^. Therefore, understanding the interactions that mediate the homing and adhesion of LT-HSC to the niche holds the potential to enhance the therapeutic efficacy of BM transplantations.

Bone marrow mesenchymal stromal cells (MSCs) are an integral component of the BM niche. MSCs produce indispensable hematopoietic factors required for proper HSC function ^2,4–6^ but also involved in the homing of hematopoietic stem and progenitor cells (HSPCs). *In vivo,* tracing of (HSPCs) revealed that upon HSPCs arrival to the perivascular niche, they establish direct, physical contact with MSCs ^7,8^. While many studies have emphasized the importance of MSCs paracrine activity, the direct cell-cell physical interactions of MSCs with LT-HSCs are much less characterized ^1,9^.

Here, we aim to investigate direct cell-cell interactions of MSCs with LT-HSCs. However, several challenges hinder the study of these interactions, limiting the experimental potential of current approaches. First, LT-HSCs are an exceedingly rare population and represent approximately 1 per 10^5^ of BM cells in mouse BM ^10–12^. Previous studies on hematopoietic cell adhesion have mainly relied on wash-based assays, which assesses cell adhesion by counting cells that remain attached to a substrate after washing^13,14^. While such assays brought valuable semiquantitative information, they usually require larger cell numbers to account for experimental variability caused by issues such as cell aggregation at the edges of culture dishes or inconsistencies in washing forces / shear rates between experiments. Next, wash-based assays do not provide detailed quantitative information about the force of adhesion and do not allow measuring early adhesion events, which occur within seconds and are critical for understanding LT-HSC homing mechanisms during the early stages^15^. Finally, BM microenvironment is rich in various types of extracellular matrix (ECM) proteins ^16,17^. Although ECM proteins play a vital role in regulating LT-HSC adhesion to the niche, their relative contributions have not been thoroughly explored. Thus, the studies on MSC and LT-HSCs interaction should also dissect the role of extracellular matrix (ECM) proteins to gain a more comprehensive understanding of the niche complexity.

To address these limitations and precisely characterize cell-ECM and cell-cell adhesion within the LT-HSC niche, we propose a novel experimental approach integrating cell sorting, surface functionalization, and atomic force microscopy (AFM) - based techniques, particularly single-cell force spectroscopy (SCFS)^18^.

Our methodology allows us to dissect the biomechanical principles governing the interactions along the ECM – stromal cells – LT-HSCs axis. We determined the contribution of the actin cytoskeleton in maintaining mechanical properties and cellular morphology of stromal cells within LT-HSCs niches and analyzed the mechanical properties of LT-HSCs isolated from young and old mice, as aging causes stiffening of bone marrow as a bulk tissue^19^. Finally, we employed SCFS to evaluate stromal cell adhesion to several niche-relevant ECM proteins and LT-HSCs adhesion to stromal cells growing on fibronectin.

The presented approach is highly adaptable and may represent an advance in the research aimed at characterizing physical interaction between LT-HSC niche components at the single-cell level.

## Results

### Actin maintains mechanical properties and morphological features of bone marrow stromal cells

Together with ECM, BM-MSCs form the scaffold of the bone marrow niche ^9^. Thus mechanical properties of bone stromal cells are crucial for maintaining the architecture of the niche. BM-MSCs are characterized by a prominent actin cytoskeleton (**Fig. 1A**, **SI: Fig S1A**) with multiple stress fibers (**SI: Fig S. 1A, white arrows**). To investigate the role of actin in maintaining the shape and the mechanical properties of BM-MSC, we used cytochalasin D (cytoD), an actin polymerization inhibitor^20^, and the model BM-MSC cell line MS-5. After cytoD treatment MS-5 was characterized by depolymerized actin cytoskeleton (**Fig. 1B**). Actin aggregation (**Fig. 1B**) was confirmed by significantly increased mean and maximal fluorescence intensities for actin (**Fig. 1. C, SI: Fig. S1B**). Cell spreading area, calculated from fluorescent images for actin cytoskeleton (**SI: Fig. S1C**), was significantly decreased after cytoD treatment (**Fig. 1D**). Additionally, to check whether the actin cytoskeleton of stromal cells is responsible for nucleus shaping, we assessed the modulation of the morphology of cell nucleus upon CytoD treatment (**SI: Fig. S2D**). After cytoD treatment, the nucleus projection area became significantly lower than in control cells (**Fig. 1E**), however nucleus to cell spread area ratio was significantly higher in CytoD treated cells, mainly due to a large drop in cell spread area (**SI: Fig. S1E**). Of note, nucleus circularity was not affected by cytoD treatment (**Fig. 1F**). To determine the contribution of actin skeleton to mechanical properties of stromal cells, we perform AFM-based elasticity measurement of control and cytoD-treated MS-5 cells, over the nuclear region of stromal cells, with bead-modified cantilevers to have a better controlled and smoother indentation (**Fig. 1G**). Stromal cells treated with cytoD remained adherent and were easily measurable with AFM (**Fig. 1H**), although changes in their morphology were visible – their lamellipodia shrank and formed longitudinal projections (**Fig. 1H, black arrow**). They were very much softer than control cells, as reflected in the change of force curve shapes (**Fig. 1I**) and their lower Young moduli. The median elastic modulus of control stromal cells was E=1243 Pa (**Fig. 1J**) and the median value of elastic modulus for cytoD-treated stromal cells was E=111.3 Pa (**Fig. 1J**). To test whether our experimental setup was stable – including the 30 minutes of incubation with cytoD and up to 45 minutes of AFM measurements – we analyzed the elastic moduli for each measured cell as a function of time. We did not observe any significant tendency in changes of elasticity values over the entire duration of micro-mechanical experiments, both for treated and untreated stromal cells (**Fig. 1K**). To test whether the geometry of the AFM tip used in elasticity measurement might affect results we further repeated this measurement with a (blunt) pyramidal tip (MLCT-DC C). While median values for both control (5534 Pa) and CytoD-treated (652 kPa) BM-MSC were around 5 times higher than those measured with spherical tip, the relative moduli difference was the same as for spherical tip (**SI: Fig S1F, G**). Previous studies also demonstrated several-fold increases in the absolute elastic modulus of samples measured with a pyramidal AFM tip compared to those measured with a spherical tip. However, the tip geometry did not affect the relative differences in the elastic moduli between the samples.^21,22^.

**Figure 1.**
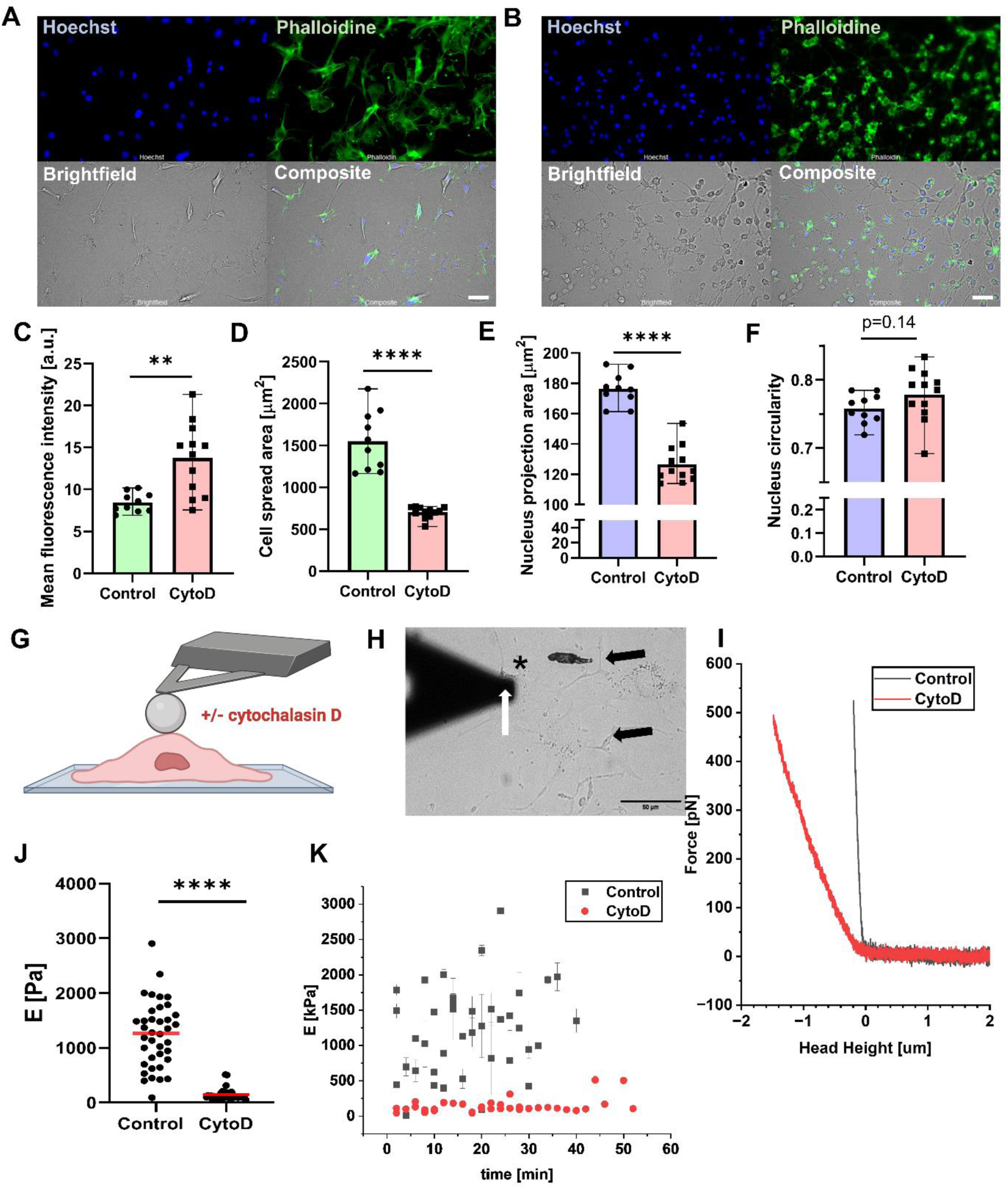
Actin cytoskeleton is responsible for MS-5 stromal cell mechanical properties. (**A**) Fluorescent images of control stromal cells (**B**) Fluorescent images of cytoD-treated stromal cells (**C**) Mean fluorescence intensity of the actin cytoskeleton. (**D**) Cell spread area. (**E**) Nucleus projection area. (**F**) Nucleus circularity. For data presented in bar plots (**C-F**) each point denotes the mean from one image on which data are obtained from: 34-89 for control and 45-242 for cytoD treated cells per image. Here **** denotes p<0.0001, ** p<0.01 for t-test. (**G**) Schematics of AFM-based characterization of stromal cells mechanical properties. (**H**) Brightfield image of experimental setup during measurement of cytoD-treated stromal cells. The white arrow denotes the spherical tip of AFM, the black asterisk the tested cell, black arrow shrank cellular projection due to cytoD action (**I**) Representative force curves recorded over control stromal cells (black) and cytoD-treated stromal cells (red) (**J**) Elastic modulus of stromal cells. Each dot presents the mean elastic modulus for one cell measured. Here, **** denotes p<0.0001 for the Mann-Whitney U test. (**K**) Scatter plot showing elastic moduli for control (black) and cytoD-treated (red) cells versus duration of the measurement.

### Stromal cells adhere rapidly to fibronectin via Arg-Gly-Asp (RGD) motive-binding integrins

Crucial stromal cells functionalities including migration, proliferation, and differentiation depend on their interaction with ECM ^16,23–27^. Therefore, in the next step, we aimed to quantify stromal cells’ adhesion to fibronectin, collagen IV, and laminin, which are crucial ECM proteins in the BM niche^16^. To achieve this, we implemented SCFS with a single stromal cell attached to an AFM cantilever positioned against a glass-bottom Petri dish coated with the respective ECM molecule as a patterned surface/surface with different zones of various composition (**Fig. 2A, SI: Fig. S2A**).

**Figure 2.**
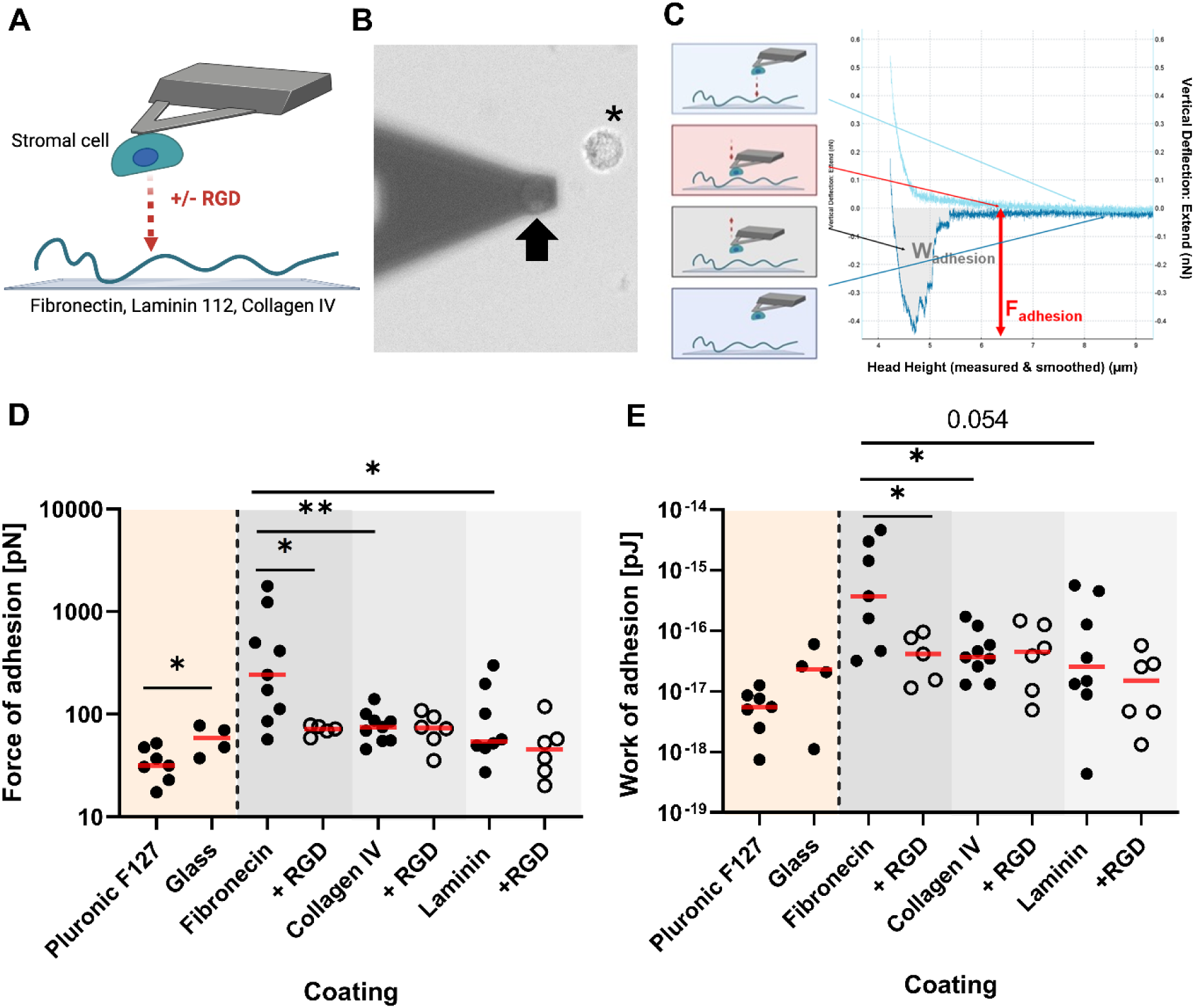
Stromal cells adhere to fibronectin via RGD-bidning integrins. (**A**) Scheme of single-cell force spectroscopy experimental setup. (**B**) Brightfield image of AFM cantilever with an attached stromal cell (thick black arrow). A floating stromal cell is also indicated (black asterisk). (**C**) Scheme illustrating each step of the single-cell force spectroscopy measurements. (**D**) Adhesion force and **(E)** work of adhesion of stromal cells to coated ECM substrates: fibronectin, collagen IV, laminin, Pluronic F-127, uncoated glass. Each dot present means for a single cell measured, open dots present means for cells measured in the presence of RGD peptideFor data presented in (**D**,**E**) red line denotes median, ***, **, * denotes p<0.001, p<0.01, and p<0.05 for Mann Whitney U test.

SCFS experiments require a first step in which cells are glued to the AFM tip functionalized with protein that ensures strong binding of the cell of interest^28^. This process can be performed either by working with non-adherent cells or by capturing detached adherent cells directly after seeding them into the dish in which the experiment is performed – before cells form tight adhesion with the substrate (**SI: Fig. S2B**). We observed that stromal cells adhere to glass and plastic very rapidly (**SI: Fig. S2C, D**), making it challenging to attach them to the AFM cantilever before they adhere to the substrate. To address this, we assessed the spreading of BM-MSC to PDMS (as a substrate) and Pluronic F127 surfactant-coated glass to determine which surface is the least adhesive for BM-MSC cells (**SI: Fig. S2C, D**). We observed that after 30 minutes, the median spread area of BM-MSC was 662.9 µm^2^ on glass, 261.4 µm^2^ on PDMS, and 177.8 µm^2^ on Pluronic F127 coated glass. After 90 minutes differences in median cell spread area were even more significant with 1544 µm^2^ on glass, 895.1 µm^2^ on PDMS, and 237.8 µm^2^ on Pluronic F127 (**SI: Fig. S2D)**. Thus, to optimize our SCFS experiments with the BM-MSC, we used homemade PDMS wells to produce patterned glass-bottom dishes. Half of the dish was coated with Pluronic F127 to facilitate BM-MSC capture onto a lectin-coated cantilever (**SI: Fig. S2E**). On the opposite side of the glass-bottom dish, we prepared three zones functionalized with different three ECM proteins of interest (**SI: Fig. S2E**). Such an approach allows us to directly compare the same cell attachment to different proteins.

Using this approach, we reproducibly attached single stromal cells to an AFM cantilever (**Fig. 2B**) and performed single-cell force spectroscopy experiments with them on ECM-coated glass (**Fig. 2C**). Passivation of glass with Pluronic F127 decreased stromal cells adhesion, reflected in a decreased median detachment force (31.57 (F127) vs 58.68 (glass) [pN]) and work of de-adhesion (2.34·10^-17^ vs 5.52·^10–18^ [J]) (**Fig. 2D,E – orange box**). Among the tested ECM proteins, stromal cells exhibited the strongest adhesion to fibronectin, as reflected by the highest median adhesion force, when compared to collagen IV and laminin, respectively (241.3 (FN) vs 74.77 (ColIV) vs 54.28 (Lam) [pN]) and work of de-adhesion (3.73·10^-16^ vs 3.67·10^-17^ vs 2.54·10^-17^ [J])(**Fig. 2D,E – grey boxes**). Importantly, in some cases, stromal cells interacting with fibronectin detached from the cantilever when pulling them away from the substrate, which indicates that the binding to fibronectin was stronger than the binding between the Concanavalin A-functionalized cantilever. This implies, that in such cases the actual force and work of adhesion are much higher than the highest measured values for similar cells on fibronectin, although, they cannot be precisely estimated (**SI: Fig. S2 F**). In conclusion, stromal cells exhibited a clear preference for adhesion to fibronectin, among the tested ECM proteins,

Given that integrin adhesion to fibronectin is often RGD-dependent ^29,30^, we tested whether blocking integrins with an RGD peptide would reduce adhesion to ECM proteins (**SI: Fig. S2G**). As expected, RGD treatment significantly decreased the adhesion of stromal cells to fibronectin with both force (241.3 (RGD+) vs 71.95 pN [J]) and work of adhesion (3.73·10^-16^ vs 4.17·10^-17^ [J]) significantly decreased (**Fig. 2D,E, grey boxes**). In contrast, RGD treatment did not affect the median force or work of adhesion for collagen IV and laminin, indicating that the adhesion of stromal cells to these substrates is independent of RGD-binding integrins (**Fig. 2D,E, grey boxes**)

### Long-term hematopoietic stem cells (LT-HSCs) are softer than bone marrow stromal cells

We showed that the mechanical properties of BM-MSC depend on the actin cytoskeleton; aide, BM-MSC adhesion to the extracellular matrix implicates the interaction between RGD-dependent integrins and fibronectin. Since bulk bone marrow mechanical properties change with aging (it becomes stiffer as showed with use of AFM^19,31^), we aimed to determine the mechanical properties of single cells: LT-HSCs derived from young and old mice and compare them to the mechanical properties of BM-MSC. We isolated LT-HSCs using fluorescent-activated cell sorting (FACS) (**SI: Fig. S3**). Mechanical characterization of cells with AFM nano-indentation requires cells to be firmly adherent to a substrate during measurement, but LT-HSCs were shown to adhere very weakly to glass^32^. Thus, we functionalized glass with concanavalin A to obtain a strong and reproducible immobilization of HSCs (**SI: Fig. S4A**). Moreover, HSCs are small cells with a reported diameter of about 5 µm^33^ (**SI: Fig. S4B,C)**, and as a consequence, we characterized their mechanics using small pyramidal tip cantilevers. The median elastic moduli obtained from cell mapping was 1.77 kPa for LT-HSC isolated from young mice, 1.26 kPa for LT-HSC isolated from old mice, and 5.53 kPa for BM-MSC (**SI: Fig. S4D**); differences between stromal cells and of both young (p=0.0011) and old (p<0.0001) HSC, were significant while both young and old LT-HSCs did not differ significantly (**SI: Fig. S3D**). This observation appears to strongly reflect the function of both types of cells. Stiff stromal cells adhere strongly to ECM giving support for soft LT-HSC, which in turn remains quiescent in their unique niche to preserve them from exhaustion and acquiring mutations.

### Single-cell force spectroscopy allows the analysis of detailed early events during LT-HSCs adhesion to bone marrow stromal cells

LT-HSCs have been shown to directly interact with stromal cells within BM niche ^8,34^, but, to date, the physical mechanisms underlying this interaction are still poorly described and understood. As a consequence, we aimed to describe LT-HSCs adhesion to stromal cells in a quantitative manner using AFM as we demonstrated for BM-MSCs. To reflect the first steps of the kinetic of LT-HSC homing to the BM niche, we measure the adhesion within seconds of MSC-HSC contact. To do so, we attached LT-HSC to cantilevers functionalized with ConcanavalinA (ConA) (**Fig. 3A**) and performed single-cell force spectroscopy experiments on stromal cells growing on fibronectin (**Fig. 3A, inset**).

**Figure 3.**
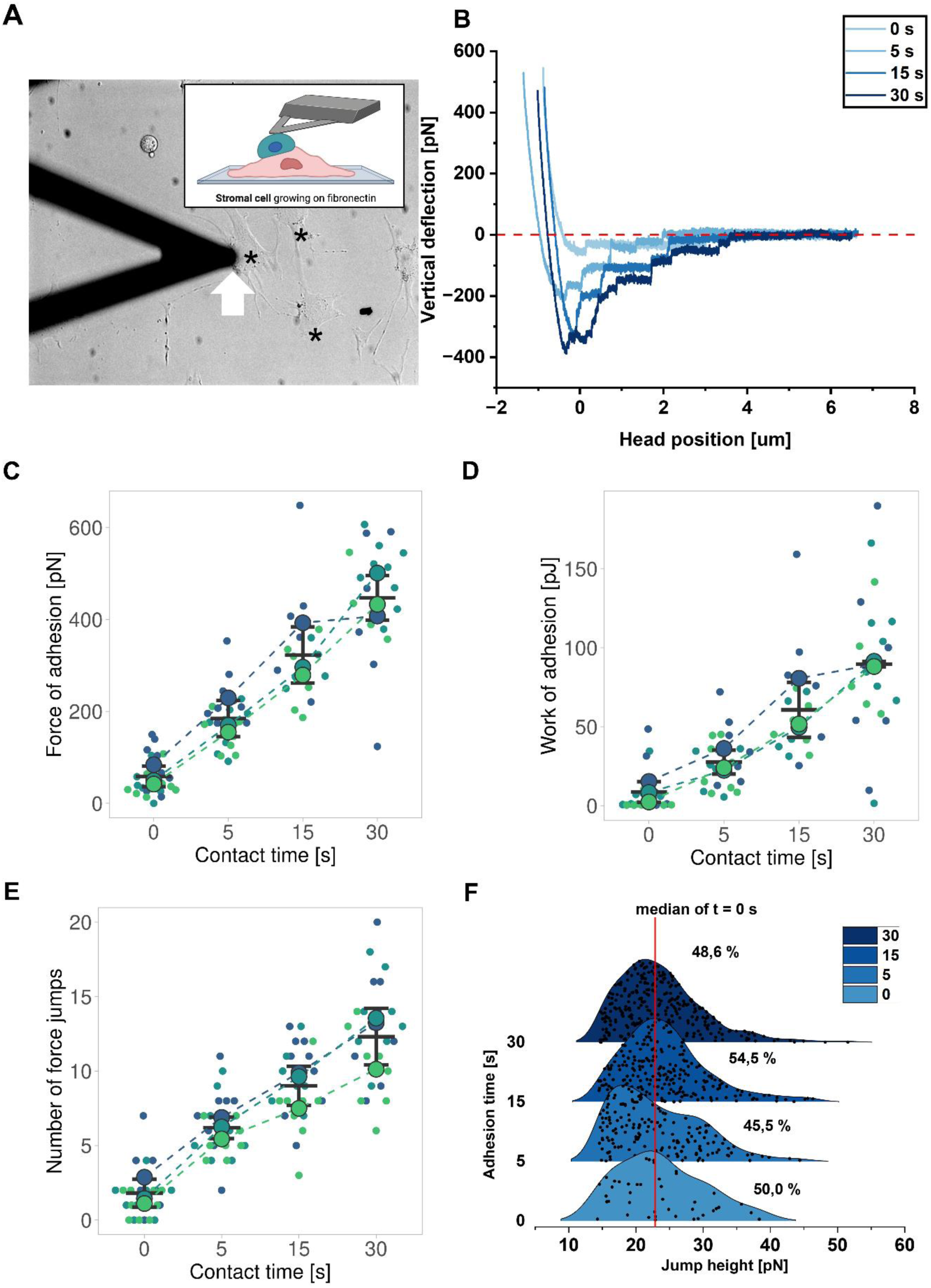
Nanoscale characterization of early events during LT-HSC adhesion to MS-5 stromal cells. (**A**) Brightfield image of AFM cantilever with a single attached long-term hematopoietic stem cell (LT-HSC) (thick white arrow). Black asterisk denotes stromal cells spread on bottom of glass dish coated with fibronectin. (**inset**) Scheme of single-cell force spectroscopy measurements. The long-term hematopoietic stem cell is attached to tipless cantilever functionalized with ConcanavalinA. Stromal cell has been growing on fibronectin. **(B)** Representative force-distance curves recorded on stromal cells with hematopoietic stem cell attached to cantilever with different cell-cell contact time. (**C**) Force and (**D**) work of adhesion of LT-HSC to stromal cell for different contact times. (**E**) Number of single-force jumps appearing in the force-distance curves. For scatter plots (**C-E**) each big framed dot represents the mean value for one stromal cell probed with LT-HSCs, small dots in the same color represent the value for a single force curve measured on the same cell, a horizontal black line denotes the mean, whiskers denote SD. (**F**) Distribution of jump heights Red line denotes median for time 0 s. The percentage of jumps with height above the median for time = 0s is depicted for each contact time.

We observed that LT-HSCs were more spread (**SI: Fig. S4E**) and less circular (**SI: Fig. S4F**) when they were attached to an AFM cantilever (**SI: Fig. S4G**), than when they were seeded directly on concanavalin coated glass (**SI: Fig. S4H**) potentially due to the stress they are subjected to during firm attachment to the cantilever. With LT-HSCs attached to the AFM tip, we performed SCFS with contact time between 0 and 30 s. We observed a gradual increase in the strength and complexity of the adhesion as a function of increasing contact time for any given cell, as reflected by force curves shape, exhibiting higher adhesion forces and more complex patterns, implicating more molecular rupture events toward full cell to substrate separation (**Fig. 3B**). Quantitatively, time-dependent strengthening of LT-HSCs adhesion to stromal cells was reflected by a rather linear increase of both mean force: 58.56 ± 22.39 (0s), 184.72 ± 39.37 (5s), 322.73 ± 61.22 (15s), 447.17 ± 48.27 (30 s) [pJ](**Fig. 3C**) and mean work of adhesion 8.65 ± 6.47 (0 s), 27.52 ± 7.54 (5s), 60.66 ± 17.34 (15 s), 89.64 ± 1.63 (30 s) [pJ](**Fig. 3D**) for contact times t = 0, 5, 15 and 30 [s] respectively.

As a complement, SCFS experiments provide not only information about “global” detachment force and work of the adhesion, but also allow us to observe the sequence of molecular events that occur during the separation of the cell from the partner substrate – namely short force jumps and longer membrane tethers (**Fig. 2C**). Those molecular events can provide more details about nature of cell-cell or cell-ECM interactions probed by SCFS, and potentially of the links of the molecules at play with the cytoskeleton^35,36^. We observed that the average number of force jumps increased with contact time: 1.81 ± 0.94 (0s), 6.19 ± 0.72 (5 s), 9.0 ± 1.31 (15 s), and 12.31 ± 1.9 (30 s) (**Fig. 3E**). We observed that the distribution of jump heights, corresponding to molecular unbinding events at single molecule or aggregate scale, does not change with time and have good stability of the median compare to zero contact time (± 5.5%) (**Fig. 3F**). Tether Plateau length distribution was shifted toward longer ones just after increasing contact time from 0 to 5 s, and remained rather similar for all contact times ≥5 s (**SI: Fig. S5A**). Nevertheless, we observed that over contact time, the population of force jumps with plateau length above 200 nm decreased from 57.1 % for 5 s contact time to around 38.7 % and 39.8 % for both 15 and 30 s contact times (**SI: Fig. S5B**). Median jump height for the population below and above 50 nm was not affected by contact time (**SI: Fig. S5C**), similarly in the case of jump populations with the cutoff of 200 nm, indicating that no collective effects of bonds are developing in this time frame, ie. that adhesion molecules are not forming laterally strong clusters that would unbind in one go at higher forces (**SI: Fig. S5C**). Thus it appears that in the first steps of LT-HSCs adhesion to stromal cells (in a time scale up to 30 s), the same type of molecules take part in the formation of adhesion sites, and over time we observe an increase of adhesion bridges between cells (**Fig. 3 E**) leading to a global strengthening of cell-cell adhesion (**Fig. 3B-D**).

Cells usually exhibiting a certain degree of spatial heterogeneity of their mechanical properties^37^ and potentially adhesive one too, we tested whether LT-HSC adhesion differs between two clearly separated and accessible to AFM zones, namely nuclear and lamellipodial compartments of stromal cells (**Fig. 4A**). Choosing a contact time 15 s, where adhesion is large enough but still allows for a full separation of the cell/cell pairs with no loss of the cell bound to the cantilever, we did not observe a significant difference in mean adhesion force (351.5 ± 109.8 pN (nucl.) vs 372.5 ± 125.5 (lam.) pN, p>0.99) (**Fig. 4B**), work of adhesion (95.4×10^-17^ J vs 62.37×10^-17^ J, p=0.4)(**Fig. 4C**) and a number of jumps (11.67 vs 9.00, p=0.69) (**Fig. 4D**) when comparing interaction of LT-HSC with nuclear and lamellipodial compartments of stromal cells. Interestingly, we nevertheless observed a significant increase in median jump forces for the lamellipodial compartment (**SI: Fig. S6A**, p=0.042), while plateau length remains unaffected by sub-cellular target localization (**SI: Fig. S6B**).

**Figure 4.**
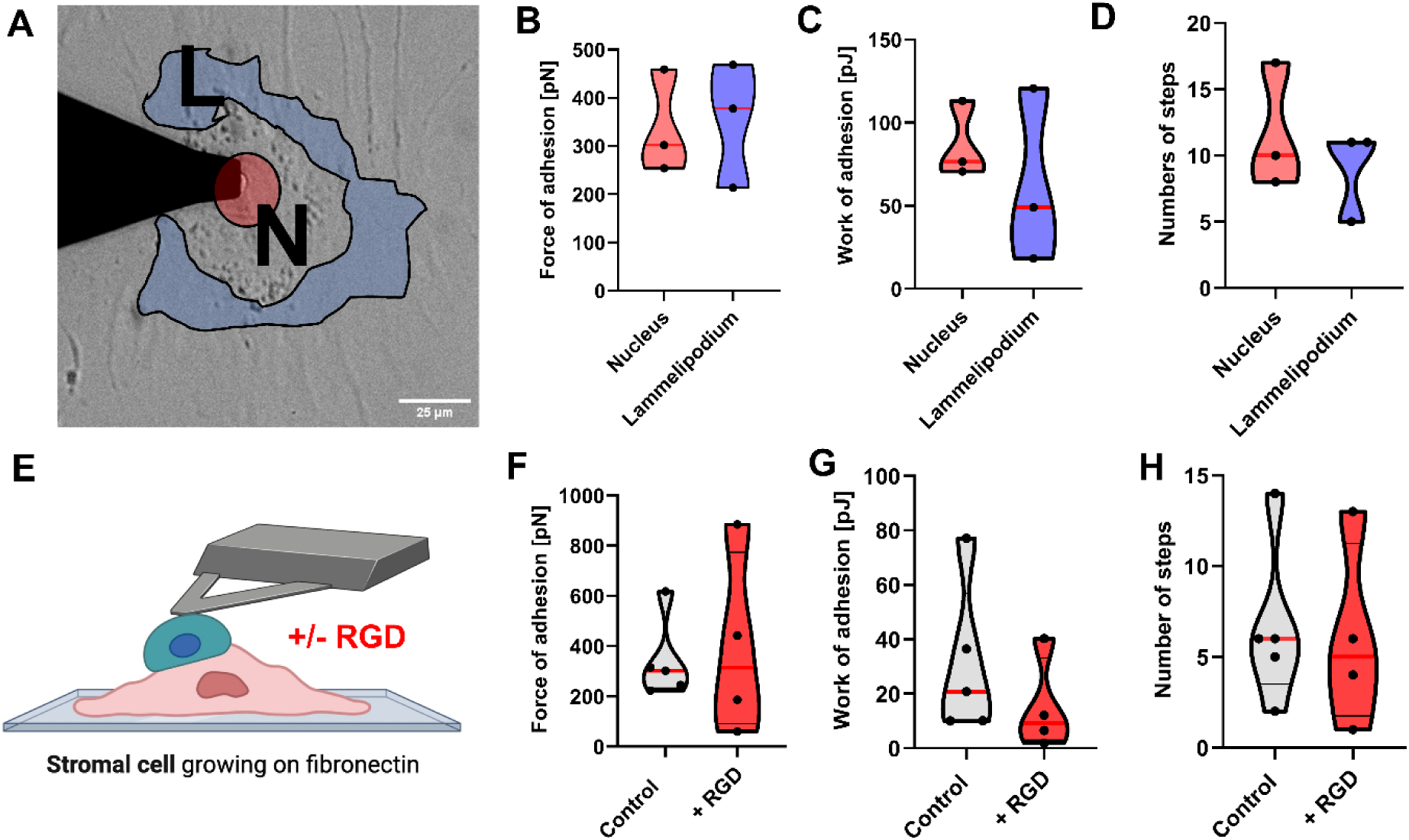
(**A**) Brightfield microscopic view of experimental setup addressing measuring LT-HSC adhesion to stromal cells above nuclear (red) and lamellipodial (blue) compartments (**B**) Force of adhesion: Each dot denotes value for single measured cell, red line denotes median. **(c)** Work of adhesion (**D**) Number of single-force jumps appearing in the force-distance curves (**E**) Schematics of experimental setup for investigating of impact of RGD on adhesion between LT-HSC and stromal cells. (**F)** Force of adhesion in presence of RGD. Each dot denotes value for single measured cell, red line detones median. (**G**) Work of adhesion (**H**) Number of single-force jumps appearing in the force-distance curves

Finally, following our demonstration that stromal cells adhere to ECM protein mainly by RGD-dependent integrins (**Fig. 2**), we questioned whether integrins might contribute to LT-HSCs adhesion to stromal cells, which could indicate a potential competition between stromal cells binding to fibronectin and LT-HSCs when homing in the niche. To do it we perform SCFS with LT-HSCs in the absence or presence of RGD (**Fig. 4E**). We do not observe significant differences between control and RGD-treated LT-HSC detaching from stromal cells, nor in the force of adhesion (356.3 ± 156.4 pN vs 401.3 ± 372.8 pN, p=0.81)(**Fig. 4F**), work of adhesion (30,2×10^-17^ J ± 27.84×10^-17^ J vs 13,23×10^-17^ J ± 16.87×10^-17^ J, p=0.43)(**SI: Fig. 4G**) nor in a number of detachment steps (3.8 ± 2.38 vs 2.75 ± 2.5, p=0.54) (**Fig. 4H**). Moreover, we do not observe significant differences between jump forces (**SI: Fig. S6C**, p=0.11) and plateau length (**SI: Fig. S6D,** p=0.14) between control and RGD conditions. We then conclude that LT-HSCs can strongly adhere within seconds after interaction with MSC and that this adhesion is not mediated by RGD-dependent integrins.

## Discussion

The aim of this study was to dissect the physical interactions between cells and ECM proteins composing the LT-HSCs niche, in particular their mechanical properties and adhesion capabilities. While multiple cell types ^4,38,39^ and ECM proteins ^16,17^ maintain LT-HSCs niche functioning, characterization of their role was up to date restricted to pure cellular and biochemical biological studies, which do not provide any quantitative descriptions of such physical parameters within LT-HSCs niche. Here, by introducing AFM-based experiments from single-cell indentation to single-cell force spectroscopy ^28,35^ we could depict quantitatively the mechanics of cells, the adhesion between stromal cells and ECM (**Fig. 2**) as well as between LT-HSCs and stromal cells (**Fig. 3,4**), providing a global biophysical description of these components of the LT-HSC cells and their niche.

LT-HSCs are at the apex of the hematopoietic system hierarchy and they are responsible for the production of all blood cells during lifespan^3^. Stromal cells are crucial for maintaining the functionality of LT-HSC niche ^2,9,40^ and generate physical contact with hematopoietic stem/progenitor cells in the process of hematopoietic stem/progenitor cells migration to the niche ^7,8^. Nevertheless, the physical mechanism responsible for adhesion between LT-HSCs and stromal cells as well as the magnitude of forces generated during adhesion were not characterized up to now. Thus in our work, we specifically focus on providing a quantitative analysis of the mechanobiology of LT-HSC and stromal cells’ physical interactions. Moreover, our approach can be further applied to analyze the interaction between different cell types with proteins, or between cells, composing the BM niche, allowing for a better understanding of mutual interaction in healthy and diseased BM.

While studies relying on time-lapse imaging of fluorescently labeled LT-HSCs niche cells^7,8^ have allowed to observation of changes in morphological features of the cells as well as their geometrical orientation in particular for their contact adhesion, such approaches cannot provide information about forces at cell/ECM or cell/cell interfaces. In contrast, atomic force microscopy, working in force spectroscopy mode, coupled to the optical observation of cells while subjected to forces, for example, to localize a cell/cell interaction at a particular place, allows to provide quantitative information about physical interactions between various samples, varying from single proteins^41,42^ up to cells^35,43,44^, or ECM constructs^45,46^. AFM force experiments can be performed in close to physiological conditions. One difficulty is that AFM-based SCFS requires immobilization of a cell of interest on the AFM cantilever and of the protein or partner cell on a substrate, which implicates verifying the innocence of the situation regarding potential mechanical biases. The attachment of a single cell requires to use of an AFM cantilever functionalized with a molecule that strongly binds the cell of interest^47^. As an example fibronectin was used to bind murine fibroblasts for SCFS^48^, as well as multiple studies included the use of mannose-binding lectin – concanavalin A^29,43,44,49^. In our work we questioned the contribution of RGD-binding integrin to the LT-HSCs vs stromal cells adhesion thus we did not use fibronectin for AFM cantilevers coating to avoid integrin-dependent activation of cells on cantilevers^50,51^. And potential detachment due to the RGD presence. Here, we observed that LT-HSCs, although reported to weakly adhere to glass^32^, adhered well to ConA-functionalized glass-bottom dishes (**SI: Fig. S4H**) as well as on AFM cantilever functionalized with ConA (**SI: Fig. S4G**). Interestingly we observed that the spreading of LT-HSCs is higher on ConA-functionalized cantilevers than on ConA-functionalized glass (**SI: Fig. S4C, E-H**). We propose to link this to the effect of pushing forces during the gluing of the very soft cells LT-HSCs to the cantilever (**SI: Fig. S2B**). This, in our case, is beneficial to maintain the LT-HSCs adherent on the cantilever SCFS in order to record cell/cell detachment forces. We observed that LT-HSCs adhesion to stromal cells increases relatively linearly with contact time (**Fig. 3D-F)**, as recently reported in other cell types^52^. For 5 s contact time between LT-HSCs and stromal cells, we detected a median adhesion force of around 200 pN (**Fig. 3C**), which can appear to be relatively low, as compared to the other cell types. Krieg et al. reported that for zebrafish embryonic ectodermal cells, the force of adhesion for 5 s contact time is around 500 pN, for endoderm around 700 pN, and for mesoderm around 600 pN. Nevertheless while comparing the diameter of embryonic cells from zebrafish which is in the range of 17-19 um, to the radius of LT-HSCs reported in our studies around 6-7 µm (**SI: Fig. S4C**), it appears that the adhesion force per area of adhesion would be similar or even higher to observed between embryonic zebrafish cells^53^. As a matter of comparison, Puech et. al observed forces in the range of 198 to 405 pN for 5 s contact time for gastrulating zebrafish cells with a mean diameter of around 17 µm^44^, Taubenberger et al. observed a mean force of adhesion of 49±7 pN for CHO cells being around 10 µm in diameter adhering to collagen matrix for 5 s^43^, finally Li et al. observed forces around 50 pN for K562 with a diameter around 20 µm adhering to fibronectin with contact time 0 s^54^. All these scales are coherent to the breaking of one to a few bonds, not largely interacting one with the other laterally (eg. as clusters or via the cytoskeleton) To better understand the mechanism of LT-HSC adhesion formation with stromal cells we analyzed the force jumps observed during LT-HSC detachment from stromal cells^43^. Such an approach allows us to dissect the contribution of particular adhesion molecules to overall adhesion – as reviewed by Helenius et al. ^35^ While we observed that the number of single detachment force jumps increased linearly with increasing contact time (**Fig. 3E**), we did not observe a shift in jump height with increasing adhesion time (**Fig. 3G**). As a contrast, Taubenberger et al. observed that CHO-A2 cells become activated after adhesion to collagen type I lasting at least 120 s, resulting in an increase in counts of higher jump heights indicating that new adhesion molecules are recruited to cell-ECM adhesion sides ^43^. In this light, we interpret our observations that in the scale of 30 s adhesion, the same type of molecules is responsible for LT-HSCs – stromal cell formation, and an increase in the force of adhesion results in the formation of higher cell-cell adhesion sites organized by the same molecules. (**Fig. 3**).

As already mentioned, bone marrow stromal cells are crucial for maintaining LT-HSCs functionality by both secreting soluble factors^2,4^ but also by providing physical anchorage for hematopoietic cells^7,8^. Here, we showed that bone marrow stromal cells’ mechanical properties are strongly regulated by a well-organized actin cytoskeleton (**Fig. 1**) as well we showed that bone marrow stromal cells are five times stiffer than LT-HSCs, regardless of the age of LT-HSCs donors (**SI: Fig. S4D**), coherent with the fact that immune cells are very often found softer than many other tissue cells^55^, in particular fibroblast^56^ or endothelial cells^57,58^. Moreover, we showed that among various ECM proteins, bone marrow stromal cells adhere stronger to the fibronectin compared to collagen IV and laminin (**Fig. 2D,E**). We further determined that stromal cells’ adhesion to the fibronectin depends on RGD-binding integrins to a great extent (**Fig. 2F,G**). This led us to question whether we can observe a competition between fibronectin and LT-HSCs as ligands for stromal cells RGD-binding integrins, which could have important consequences for the LT-HSCs niche organization (**Fig. 4E**). Interestingly, we did not observe a significant impact of RGD on LT-HSCs adhesion to stromal cell (**Fig. 4 E-H)**, in line with previous results showing that RGD-binding integrins are crucial for cellular adhesion to ECM proteins. ^44,49,49,59^ but are less implicated in cell-cell adhesion formation^60,61^. On the other hand, while stromal cells are adhering to fibronectin, all RGD-binding integrins expressed by stromal cells might be engaged in the formation of adhesion with the ECM present on the basal surface (coated glass) potentially limiting their interactions with LT-HSCs via intercellularly present, potentially secreted, ECM.

To conclude, we showed that AFM has the capability to quantitatively characterize physical interaction within the LT-HSC niche. Such single-cell techniques appear to be, in our hands, well suited to the biophysical studies of rare, small, and delicate cells. With our approach, we are capable of determining the mechanical properties of various niche-building cells and precisely describing the adhesion of a given cell type within a niche even for rare cells since SCFS uses one cell at the time to probe surfaces or other cells. We are capable of deciphering the contribution of particular adhesion molecules into the adhesion process (**Fig. 2F,G**, **Fig. 4F-H**) but also recording precisely depict the time-dependent dynamics of cell-cell adhesion organization in a time-dependent manner (**Fig. 3, SI: Fig S5,6**). This opens a new direction in sequentially analyzing the interactions within the bone marrow niche. We foresee that the presented approach may help to decipher the adhesion molecules which determine the specificity of LT-HSCs homing and adhesion to the niche in the bone marrow. Finally, the possibility to prospectively block selective interactions during the measurements may help to screen for new specific LT-HSC drugs that mobilize LT-HSC from BM to peripheral blood.

## Materials and methods

### Surface coating ECM

To pattern the glass surface for cell culture with ECM, we used polydimethylsiloxane (PDMS) wells. PDMS was prepared from SYLGARD® 184 SILICONE ELASTOMER KIT in a ratio of 10:1 base/curing agent according to manufacturer instructions. By using a biopsy punch or a scalpel, we cut wells in the PDMS block to set the surface for a particular experiment (**SI: Fig. S2A**). PDMS blocks with wells were sterilized by putting and stirring them into 70 % ethanol. Before use, PDMS blocks were dried on a Petri dish cover under a gas run to evaporate the remaining ethanol from PDMS. PDMS blocks were placed on the sterile Petri dish glass surface with the use of tweezers and gently pressed to generate firm adhesion. An appropriate volume of 0.1 mg/mL solution of ECM proteins (fibronectin, collagen IV, laminin) in 0.1 M Tris-HCl (ph=7.4) was inserted in each PDMS wells (i.e. for wells on **SI: Fig. S2A** around 10 µL per well). Coating lasts 1 hour at room temperature, protected from dust. After this time protein solutions were aspirated, the wells washed and PDMS was removed with the use of a tweezer and the Petri dish was washed twice with PBS.

### Fabrication of bead-decorated AFM cantilevers

To prepare AFM cantilevers with spherical tips, we used tipless MLCT-O C cantilevers (Bruker). Polystyrene beads (diameter of 9.6 µm, Invitrogen) in suspension in MQ water to remove salts were spread and dried on a glass slide to obtain separated single beads. We deposited transparent 2-component epoxy glue (SADER, 5 min drying time) near the spread beads. Then we gently dip the lever tip in epoxy glue, remove if needed the excess of glue, and then catch a single spherical probe with the cantilever. Once the probe was attached to the cantilever micromanipulator, it was withdrawn and left untouched for 5 minutes. After this time cantilever was taken out from the micromanipulator and stored for at least a day at room temperature before use to ensure final proper glue hardening.

### Cell culture

Culture of bone marrow stromal cells (MS5) was performed in αMEM cell culture media (Gibco) supplemented with 10% fetal bovine serum (Biowest) and 1x Pen-strep. MS-5 cells were passaged 2-3 times a week in a ratio of 1:6 to avoid over-confluence. Before passage, cells were washed with PBS, harvested with the use of TryplE (Gibco), and centrifugated with 400 RCF for 5 minutes. After passage cell number was determined with the use of a counting chamber (Bürker and Malassez).

### Cytochalasin D treatment

A stock solution of Cytochalasin D (CytoD, Sigma) was prepared in a concentration of 10 mM in DMSO, it allowed us to the final 10 µM CytoD concentration in a cell culture medium, ensuring that DMSO diluted 1000x not affect the biomechanical properties of cells ^62^.

### Immunofluorescent staining

For immunofluorescence staining, cells were seeded on a glass dish coated with 0.1 mg/ml fibronectin. After 24 hours in culture condition, the cells were fixed with 4% paraformaldehyde for 20-30 minutes. The cell membrane was permeabilized by treatment with cold, 0.2 % Triton X-100 solution for 5 minutes. Subsequently, blocked with 0.5% BSA for 1 hour and then stained with Alexa488-conjugated Phalloidin (1:200) for 1 hour to visualize actin. To visualize cell nuclei, cells were stained with Hoechst 33342 (1:5000) for 15 minutes, before washing the cells. Both staining steps were performed at room temperature.

### Fluorescent imaging

For fluorescent imaging, we used a fluorescent microscope Axio Observer (Zeiss) equipped with Apotome 3 (Zeiss), Axiocam 702 mono camera (Zeiss), and multicolor led source Colibri 7 (Zeiss). Images were acquired with 20x/0.8 DICII Plan Apo air objective. The imaging system was controlled by ZEN 3.6 (Zeiss) software. All quantitative image analyses were done with the open-source software Fiji^63^. For confocal imaging of stromal cells actin cytoskeleton, we used Leica Stellaris. For Phalloidin imaging we used a 488nm (20 mW) laser. Images were acquired with the use of 63x PlanApo objective with oil immersion. The confocal image postprocessing was done with the use of LasX software (Leica).

### Image analysis

Image analysis was performed using open-source Fiji Imagej software as previously described^64^. All fluorescent images for the given channel used for comparison were recorded at the same imaging parameters(laser, intensity, exposure time, and threshold). Schematic presentation of image processing is presented in **SI: Fig. S1C, D**. Image type was set to 8bit, and a threshold was set to cover the cell spread area or nuclear projection area (**SI: Fig. S1C, D**). For nuclei images, particle analysis was performed after applying a threshold for the area covered by nucleus projection to obtain information about nuclei number, nucleus projection area and nuclear circularity. The mean cell spread area for each image was then calculated as follows:

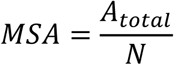

Where MSA denotes mean spread area per cell, *A_total_* denotes area covered by cells, and *N* denotes nuclei number. Finally for 8-bit images mean fluorescent intensity was determined from grey value histograms.

### Animals

C57BL6 mice were obtained from The Jackson Laboratory and housed under specific pathogen-free conditions in the animal facility at the Faculty of Biochemistry, Biophysics, and Biotechnology of the Jagiellonian University in Cracow and at Centre d’Immunologie de Marseille-Luminy (CIML). All animal procedures and experiments were performed in accordance with the limiting principles for using animal in testing (the three Rs: replacement, reduction, and refinement) and approved by the French Ministry of Higher Education and Research or in accordance with the guidelines set forth by the NIH Animal Care and Use Committee. Animal experiments were done in accordance with institutional animal care and ethical committees and Polish, French and European guidelines for animal care.

### Hematopoietic stem cell isolation

LT-HSCs were isolated as described with details in: ^65,66^. Briefly, tibias and femurs were dissected, and crushed with mortar and pestle in FACS buffer (2% fetal bovine serum (FBS) in PBS with 100 U/ml DNase), and the supernatant was collected. Then cells were negatively isolated using a Lineage Cell Depletion Kit, mouse (Miltenyi Biotec), and LS MACS columns (Miltenyi Biotec), according to the manufacturer’s instruction.

### Sample staining and cell sorting

Fraction of BM cells negative for lineage markers were stained for 30 min with DAPI for viability assessment and a cocktail of directly conjugated antibodies: lineage markers (CD3, Gr-1, CD11B, B220, and TER119), c-Kit, sca-1, CD48, CD150, CD34. Catalog numbers, concentrations, and clone information are provided in **SI: Table 1**.

Cell sorting was done on BigFoot Spectral Cell Sorter (Thermo Fisher Scientific). Singlets were selected based on FCS-A vs. FCS-W and SSC-A vs. SSC-W. Dead cells were excluded. Upon live cells HSCs were defined as Lin^-^ c-kit^+^ sca-1^+^ CD48^-^ CD150^+^ CD34^-^. The gating strategy is shown in **SI: Fig. S3A**.

### Determination of cell elasticity using AFM

To determine cell elasticity we used two AFMs equipped with JPK Nanowizard 1 and 2 heads. Photodetector sensitivity was determined by firstly recording the calibration force curve on the glass substrate and determining the inverted slope on the extended force curve in JPK software. The spring constant of each cantilever was determined by acquiring thermal noise and determining the resonance frequency following manufacturer methodology. In experiments with the use of a spherical AFM tip, 5 force curves per cell were acquired over the nuclear region of stromal cells with contact F = 500 pN and displacement v = 8µm/s. In experiments with the use of MLCT-C BIO cantilevers (Bruker), elasticity maps with sizes of 5 x 5 and 8 x 8 px were acquired with contact F = 700 pN and displacement v = 8µm/s. Hertz-Sneddon model was then fitted in JPK Data Processing software in order to determine the elastic modulus of cells (denoted Young modulus).

### AFM cantilevers functionalization with ConcanavalinA

To firmly attach cells to cantilevers their functionalization was performed like previously established ^47^. Compounds used are presented in **SI: Table 2**. Briefly, 1 min residual air plasma-cleaned MLCT-O cantilevers were incubated overnight with 0.5 mg/mL biotinylated-BSA at 4°C. After overnight incubation, cantilevers were washed three times in PBS and incubated in 0.5 mg/mL solution of streptavidin for 30 minutes at room temperature. Cantilevers were washed three times with PBS and incubated for 30 minutes at RT in 0.4 mg/mL solution of biotinylated ConA solution, followed by three times washing in PBS. Functionalized cantilevers were than stored in PBS at 4°C and used within 10 days after functionalization.

Functionalization of glass coverslips with lectins for measurements of LT-HSC elasticity was performed with the same protocol as for AFM cantilevers for SCFS.

### Single-cell force spectroscopy

For SCFS experiments Fluorodish coated with ECM and Pluronic F127 were used. For determination of stromal cells adhesion to ECM Fluorodish with coating pattern presented in **SI: Fig. S2E** was applied, while for probing adhesion of LT-HSC to stromal cells, only stromal cells growing on fibronectin-coated glass were used. Each cantilever was calibrated before measurement in the same manner as described previously, before catching a cell. Cells suspensions were pipetted over the Pluronic F127 coated region. We attached cells to ConcanavalinA-functionalized cantilevers by pressing a non adherent cell with the cantilever onto the non adherent F127 surface (**SI: Fig. S2B**), retract the lever away from the surface, wait 5 minutes for the cell to get firmly attached to the cantilever (**Fig. 2B, SI: Fig. S4G**) and then performed single-cell force spectroscopy (**Fig. 2C**) over zones of the petri dish coated with a particular ECM protein and/or over stromal cells growing on fibronectin. After this time force curves were recorded over a particular ECM with contact F = 500 pN and displacement v = 2 µm/s. We set the contact condition to be at a constant height not to press continuously on the cells during the desired duration (0 to 30 sec). Adhesion force, work of adhesion, and the number of jumps were determined with JPK Data Processing software.

### Statistical analysis

For statistical analysis and data visualization, we used GraphPad Prims and Origin2022b software. Before samples were compared, the normality of the distribution was evaluated with the Shapiro-Wilk test. If all samples chosen for comparison were found normally distributed, we used the t-test for comparison. If the normality of distribution for one or both samples was rejected, we used a Whitney U Mann nonparametric test. We applied the following system of significance for all data presented in this manuscript: **** - p<0.0001, *** - p<0.001, ** p<0.01 and * - p<0.05.

## Declarations

### Ethics approval and consent to participate

All animal procedures and experiments were performed in accordance with national and European legislation.

### Consent for publication

Not applicable.

### Availability of data and materials

The data are available in the presented manuscript and supporting files. Raw data (force curves maps in format .jpk-force-map) and microscopic images (in format either .tif or .czi) are available from corresponding authors upon reasonable request.

## Authors contributions

AK, PHP, and KS designed research. AK, NB-K., KK, KG, PK, PKo, MBP, JH, TN, ME, MD, OL, M.T-K.; performed research; AK, NBK., PK, ME, analyzed data, AK, PHP and KS interpreted data, AK, NBK, KG – prepared figures, AK, NBK, PK, PHP and KS wrote the manuscript. AK, NBK, KK, KG, PK, MBP, PHP, and KS edited manuscript. All authors approved the final version of the manuscript.

## Acknowledgments

Purified human fibronectin was a gift from Peter McCourt (University of Tromsø, Norway). FYI it was purified from human plasma using Cytiva’s gelatin-Sepharose according to their instructions. Authors acknowledge the support of Beata Rysiewicz from Laboratories of Multi-Level Imaging of Biological Structures (Jagiellonian University) in the acquisition of confocal fluorescent images of stromal cell. Parts of the figures were created using <u>BioRender.com</u>.

