## Supplementary information for "AFM-based nanoscale characterization of physical interaction within hematopoietic stem cells niche at single-cell level"

### Supplementary info: Supplementary figures caption

**Supplementary Figure 1**. (**A**) Confocal image of the actin cytoskeleton of MS-5 stromal cells stained with phalloidin. The white arrow indicates stress fibers. (**B**) Maximal fluorescence intensity for actin cytoskeleton 8-bit images. (**C**) Schematic of data analysis for fluorescent images of actin cytoskeleton. (**D**) Schematic of data analysis for fluorescent images of cell nuclei. Count, area, and shape descriptors were determined for each masked set of nuclei (**E**) N/C ration – nucleus projection area/cell spread area value. For data presented in bar plots (**B**,**E**) each point denotes the mean from one image on which data are obtained from: 34-89 cells per image for control and 45-242 cells per image for cytoD-treated cells. Bar denotes mean. **** and ** denotes $p<0.0001$ and $p<0.01$ in t test. (**F**) Distribution of elastic moduli for control (black line) and cytoD (red line) treated stromal cells(with spherical tip). (**G**) Distribution of elastic moduli for control (black line) and cytoD (red line) treated stromal cells (with pyramidal tip).

**Supplementary figure 2** (**A**) Photography of experimental setup used for coating of glass with ECM proteins. PDMS with wells was attached to fluorodish and each well was filled with ECM solution of interest. (**B**) cell attachment procedure to cantilever functionalized with ConA (**C**) Brightfield microscopy images of stromal cells during their spreading on different substrates times 30 and 90 minutes. (**D**) Cell spread area on the different substrates at times 30 and 90 minutes. Each dot presents area of one cell, red line median. (**E**) Coating of glass-bottom Petri dish **(F)** Line+symbol plot showing percentage of cells attached to cantilever during constitutive force-ramps up to 5 in each single-cell force spectroscopy measurements for ECM-coated and control glass. (**G**) Experiment with RGD blockade of integrins. Left force curve is without RGD peptide in culture medium, right force curve is after incubation with 250 μM RGD peptide.

**Supplementary figure 3** Gating strategy for isolation LT-HSC from murine bone marrow.

**Supplementary figure 4** (**A**) Scheme of experimental setup. A given LT-HSC is immobilized on the glass by concanavalin A. (**B**) Topography map of single LT-HSC. (**C**) LT-HSCs diameter calculated from brightfield images of LT-HSCs adhering to glass compared to spreaded on AFM cantilevers. (**D**) Distribution of median moduli determined for single cells fitted with Kernel Smooth for LT-HSC from young (2 month), old (24 month) and for MS-5 stromal cells. Dashed lines denotes median value for each cell type. (**E**) LT-HSCs cell spread area calculated from brightfield images of LT-HSCs adhering to glass and spreaded on AFM cantilevers. (**F**) LT-HSCs circularity was calculated from brightfield images of LT-HSCs adhering to glass and spread on AFM cantilevers. (**G**) Representative image of LT-HSCs attached and spread on AFM cantilever, arrow indicates cell location. (**H**) Brightfield image of LT-HSCs adhering to ConA-functionalzied glassbottom cell culture dish, arrows indicate LT-HSCs.

**Supplementary Figure 5** (**A**) Distribution of plateau length for jumps determined for adhesion between LT-HSC and MS-5 stromal cells for contact time t= 0s , t=5s, t=15s, t=30s. Red line denotes median for time 0 s, percentage of jumps with plateau length above is depicted for each contact time. (**B**) Scatterplots showing relationship between the height of jumps and plateau lenght for each analyzed jump. Red line denotes the value of plateau length of value 50 nm. (**B**) Distribution of force jump heights for population with plateau lenght below and above 50 nm. Median is denoted as red line. (**C**) Distribution of force jump heights for population with plateau lenght below and above 200 nm. Median is denoted as red line.

**Supplementary figure 6**. (**A**) Distribution of jump heights for adhesion between LT-HSCs and MS-5 stromal cells for nuclear and lamellipodial compartments. The red line denotes the median for the nuclear compartment. The percentage of jumps with height above the median for nuclear region is shown. (**B**) Distribution of plateau length for adhesion between

LT-HSCs and MS-5 stromal cells for nuclear and lamellipodial compartments. The red line denotes the median for the nuclear compartment. The percentage of plateau length above the median for the nuclear region is shown. (**C**) Impact of RGD treatment on the distribution of jump heights for adhesion between LT-HSCs and MS-5 stromal cells. The red line denotes the median for the control condition. The percentage of jumps with height above the median for the control condtition is shown. (**D**) Impact of RGD treatment on the distribution of plateau length for adhesion between LT-HSCs and MS-5 stromal cells The red line denotes the median for the control condition. The percentage of plateau length above the median for the control condition is shown. (**C**) Distribution of jump heights for adhesion (**D**) Distribution of plateau length for adhesion .

**Supplementary info: Supplementary Figure 1.**

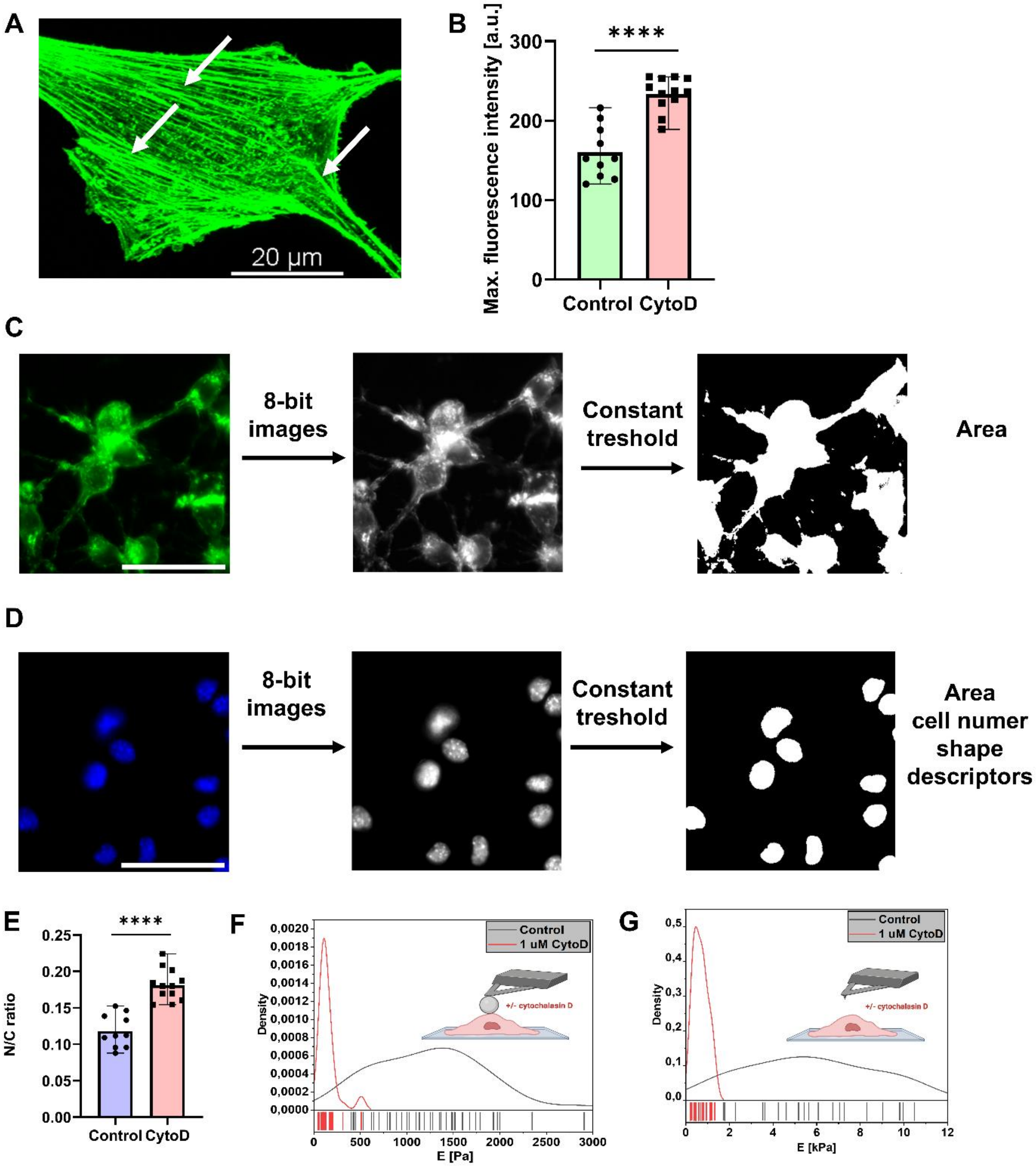

**Supplementary information: Supplementary Figure 2.**

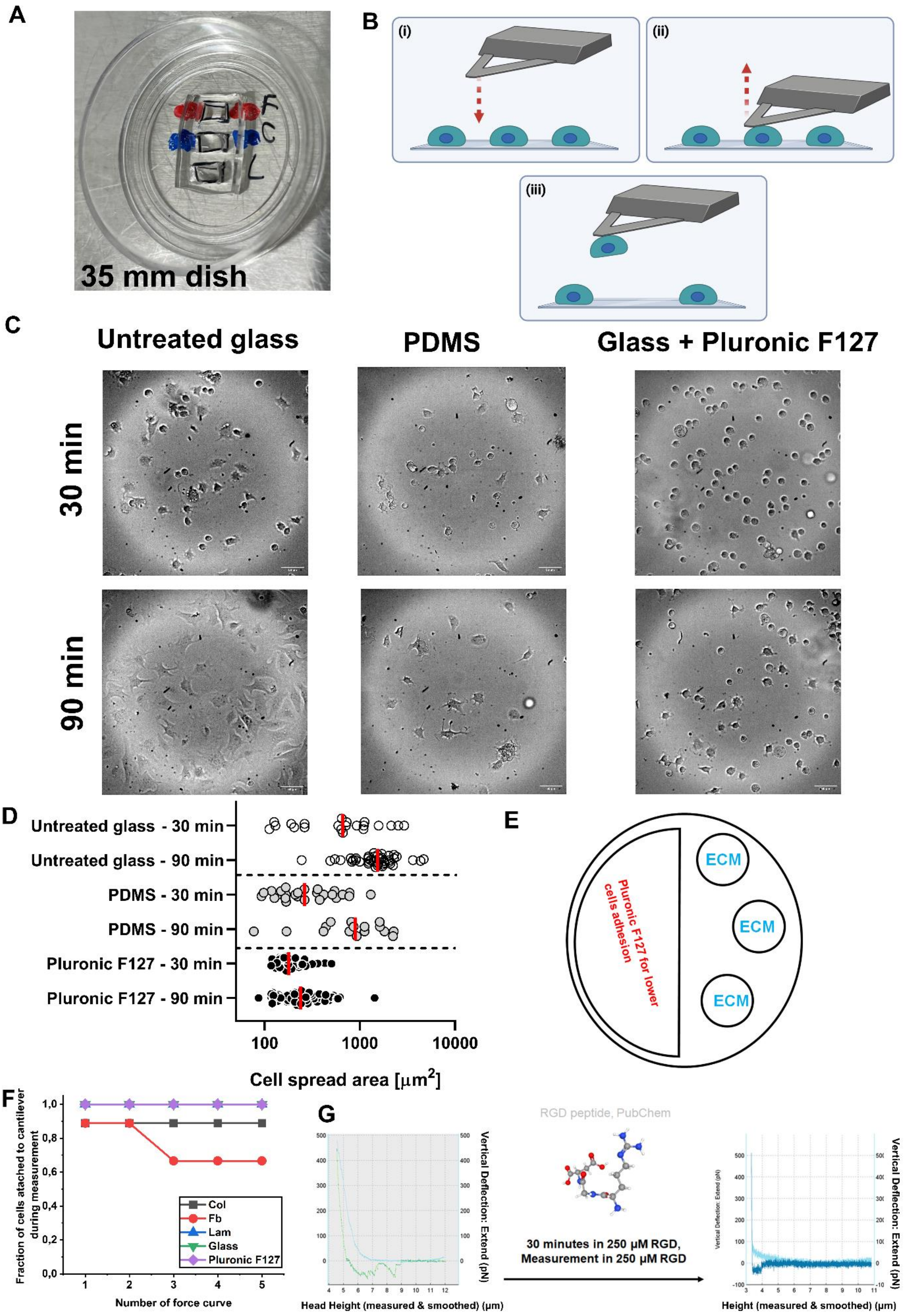

**Supplementary information: Supplementary Figure 3.**

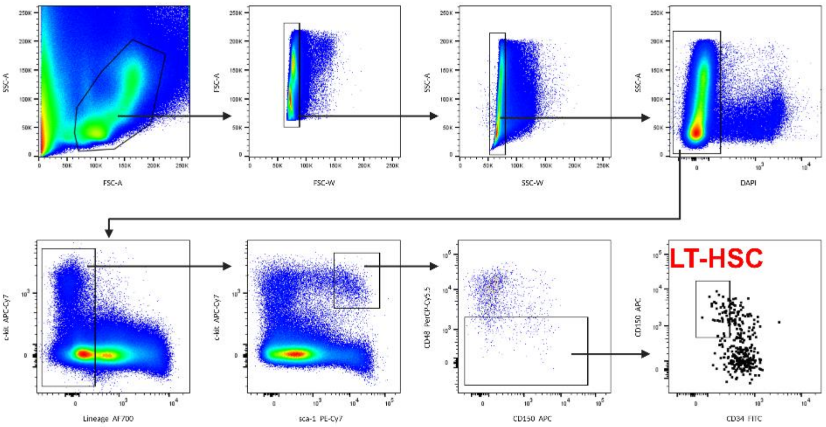

**Supplementary information: Supplementary Figure 4.**

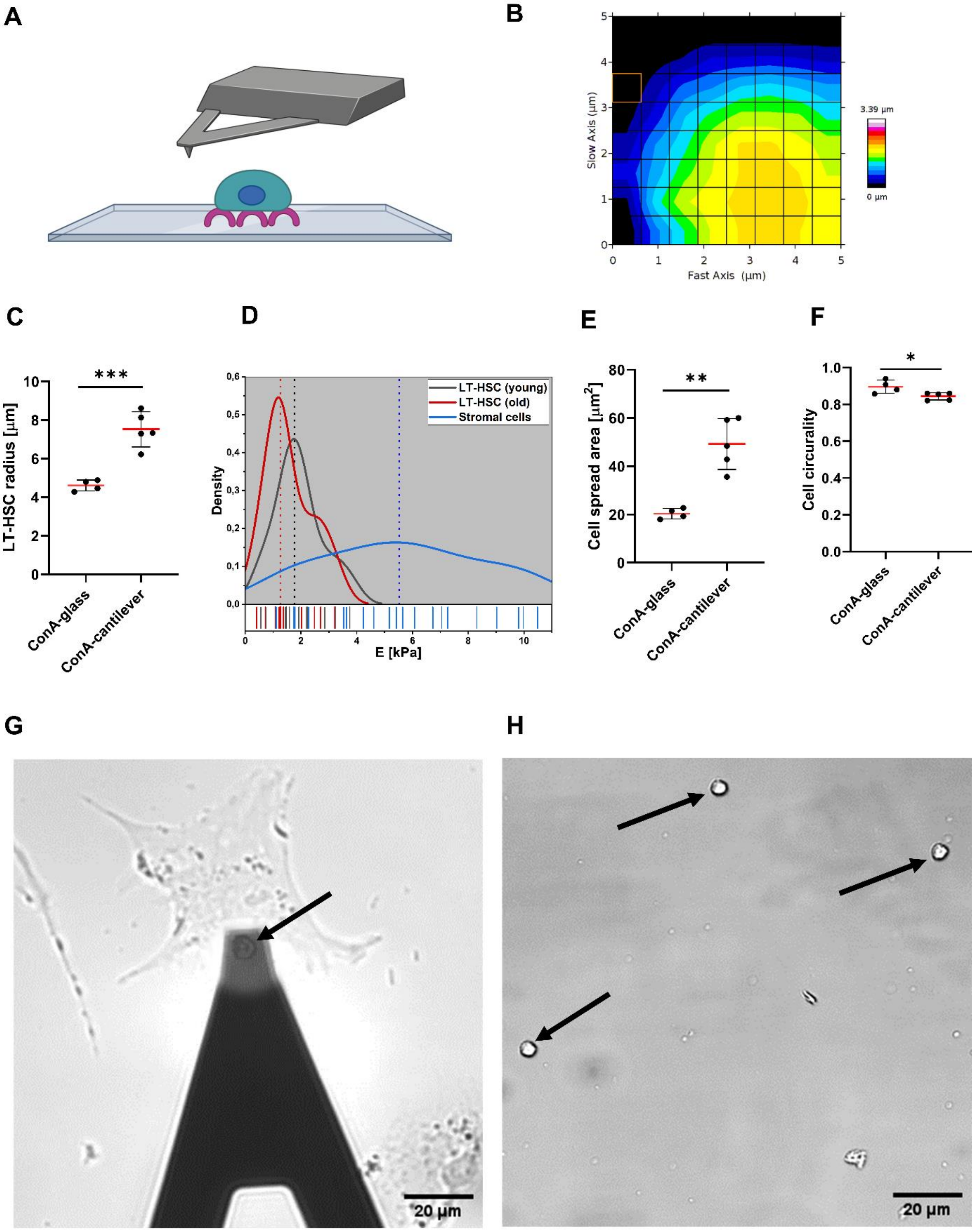

**Supplementary information: Supplementary Figure 5.**

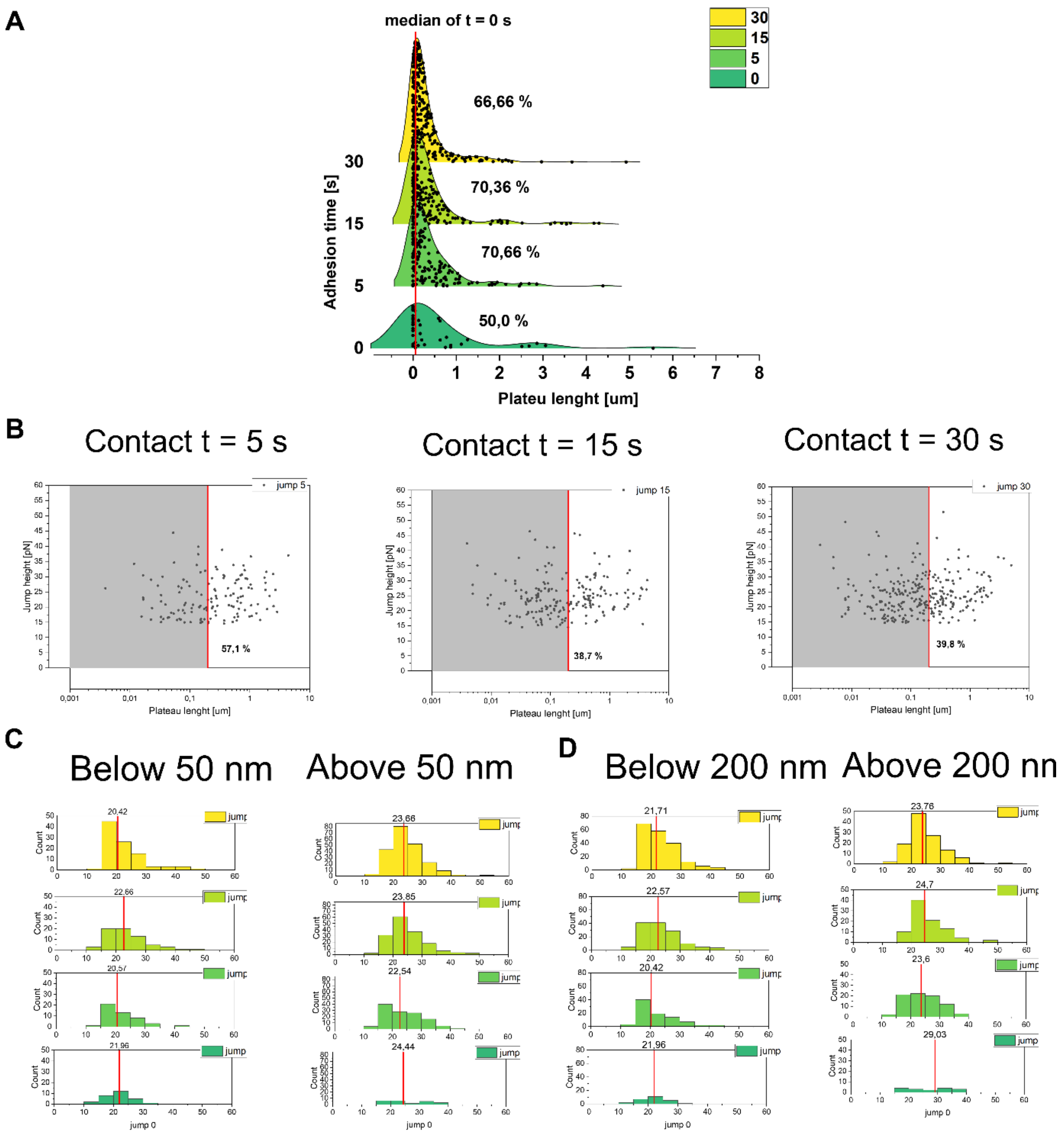

**Supplementary information: Supplementary Figure 6.**

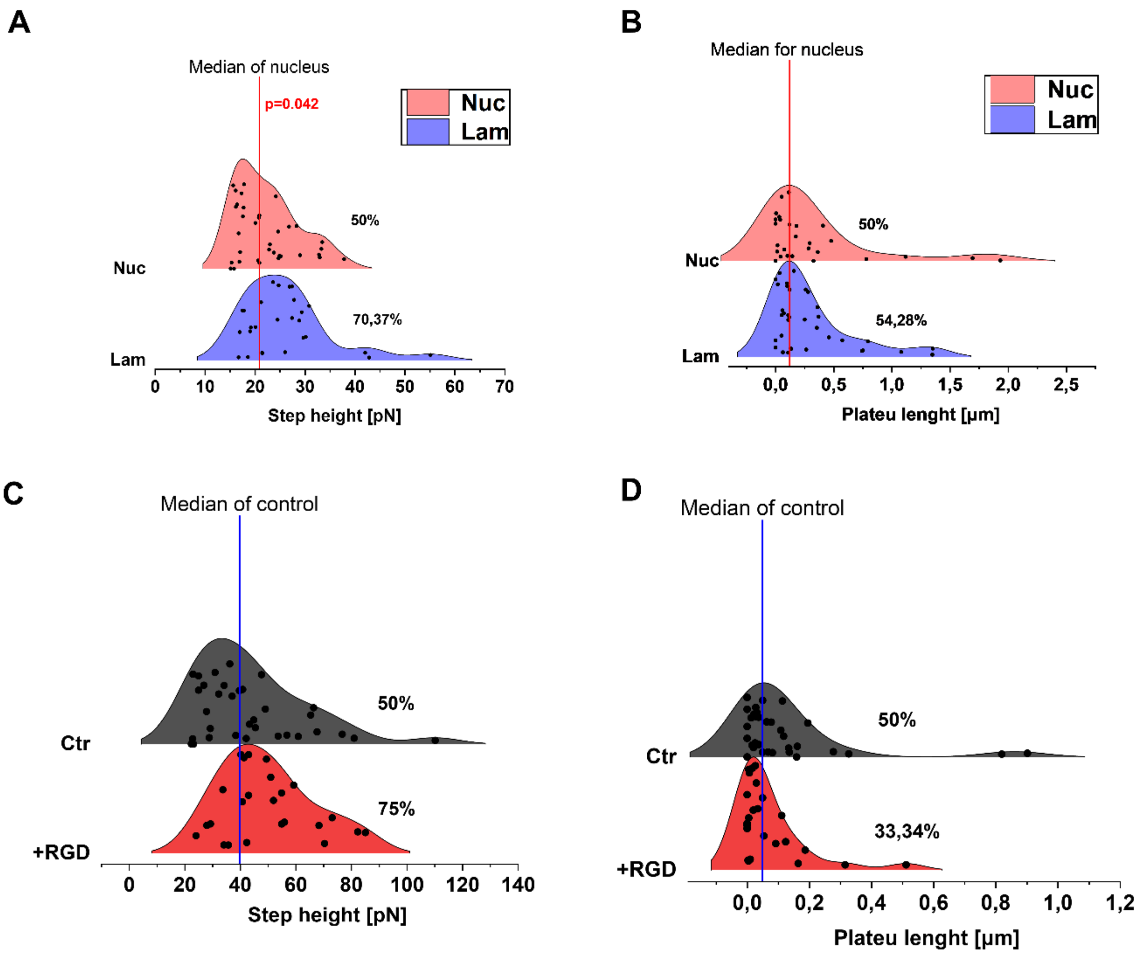

**Supplementary information: Supplementary Table 1**

Antibodies used for staining of murine bone marrow for LT-HSCs isolation with FACS

| Marker | Clone | Fluorophore | Catalog number | Concentration |
|---|---|---|---|---|
| lineage:<br>CD3<br>Gr-1<br>CD11b<br>B220<br>Ter119 | <br>17A2<br>RB6-8C5<br>M1/70<br>RA3-6B2<br>Ter-119 | AF700 | BioLegend 133313 | 1:20 |
| c-Kit | 2B8 | APC-eFluor780 | eBioscience 47-1171-82 | 1:50 |
| sca-1 | D7 | PE-Cy7 | BioLegend 108114 | 1:50 |
| CD48 | HM48-1 | PerCP-Cy5.5 | BioLegend 103422 | 1:50 |
| CD150 | TC15-12F12.2 | AF647 | BioLegend 115918 | 1:50 |
| CD34 | RAM34 | FITC | eBioscience 11-0341-85 | 1:50 |

**Supplementary information: Supplementary Table 2**

Reagents for functionalization of cantilever with ConA and times for each step of incubation.

| **Name** | **Stock concentration** | **Solvent** | **Working solution** | **Solvent** | **Time of incubation in 50 uL droplets** |
|---|---|---|---|---|---|
| Biotinylated-BSA | 2 mg/mL | Disiled $H_2O$ | 0.5 mg/mL | 100 mM pH 8.6 NaHCO3 buffer (adjusted with 1 M NaOH) | Overnight in (37C humified) |
| Streptavidin | 1 mg/mL | PBS (w/o Ca2+, Mg2+) | 0.5 mg/mL | PBS (w/o Ca2+, Mg2+) | 30 min, RT |
| Biotinylated-ConcanavalinA | 0.8 mg/mL | PBS (w/o Ca2+, Mg2+) | 0.4 mg/mL | PBS (w/o Ca2+, Mg2+) | 30 min, RT |
